# Timing along the cardiac cycle modulates neural signals of reward-based learning

**DOI:** 10.1101/2022.07.07.498947

**Authors:** Elsa Fouragnan, Billy Hosking, Yin Cheung, Brooke Prakash, Matthew Rushworth, Alejandra Sel

**Author notes:** Authors contributed equally to this work (Matthew Rushworth, Alejandra Sel).

## Abstract

Natural fluctuations in cardiac activity influence brain activity associated with sensory stimuli and affect perceptual decisions about low magnitude, near-threshold stimuli. However, little is known about the impact of fluctuations in heart activity on other internal representations. Here we investigate cardiac influences on learning-related internal representations – absolute and signed prediction errors. By combining machine learning techniques with electroencephalography (EEG) and both simple, direct indices of task performance and computational model-derived indices of learning, we demonstrate that just as people are more sensitive to low magnitude, near threshold sensory stimuli in certain cardiac phases, so are they more sensitive to low magnitude absolute prediction errors in the same cycles. Importantly, however, this occurs even when the low magnitude prediction errors are associated with clearly suprathreshold sensory events. In addition, participants exhibiting stronger difference in their prediction errors representations between cardiac cycles exhibited higher learning rates and greater task accuracy.

## Introduction

In situations where we must make decisions based on noisy or incomplete information - for example deciding whether to cross the street on a foggy morning with poor visibility - the timing of the cardiac cycle can modulate our perception by facilitating or impeding stimulus detection. Studies investigating near-threshold sensory events, like visual, auditory or somatosensory events, have shown that timing in the cardiac cycle (e.g. whether events happen during the systolic or diastolic phases of the cardiac cycle) impacts the perception of sensory cues through changes in associated neural signals^1,2^. Although heart-brain interactions are starting to be understood in relation to sensory-driven processes, it is unclear whether the cardiac cycle has a similar impact on other internal representations which are non-sensory but which, like sensory stimuli, shape decision making. Here we focus on a much-studied internal representation – the reward prediction error [PE] – and investigate whether the cardiac cycle also determines the impact that each PE will have on learning. Importantly, the magnitude of the PE can be dissociated from the magnitude of the accompanying sensory stimulus. This makes it possible to determine whether the cardiac cycle has an impact on near-threshold PEs even if the PEs are associated with clearly suprathreshold sensory stimuli.

Adaptive decisions rely on accurate subjective value estimates associated with past experience of choices and their consequences. These values can be formally defined through the reinforcement learning framework^3^ that uses the difference between expectation and outcome (the PE) to update values associated with choices. A choice that led to a positive outcome is more likely to be associated with a higher value than a choice that did not. Within this framework, learning is driven by two separate outcome dimensions: the signed PE, representing how much better or worse the value of an outcome is compared to what was expected, and the absolute PE (also called ‘salience’ or ‘surprise’), representing how much an outcome differs from expectations regardless of whether it is better or worse. Whereas positive and negative signed PEs lead to the reinforcement or extinction of the choices that led to them^4^, the absolute PEs determine the extent to which the associations between outcome and expectations need to be adjusted^5,6^. Even if a choice leads to a clearly suprathreshold sensory event, the PE it entails might be large, small, or even near-threshold depending on what the decision maker’s prior expectations were. This means that we can examine whether near-threshold PEs are impacted by the cardiac cycle even if they are associated with suprathreshold sensory events.

The cardiac cycle is a series of contractions and relaxations that help the heart pump blood throughout the body. Each cardiac cycle has a diastolic phase (also known as diastole) in which the heart chambers relax and fill with blood, and a systolic phase (also known as systole) in which the heart chambers contract and pump blood to the periphery. These two physiological phases are differentially signalled to the brain through baroreceptor firing during systole and by a pause in firing during diastole. These signals are linked to activity in brainstem regions such as periaqueductal gray as well as forebrain regions such anterior cingulate cortex (ACC), anterior insula (AI), amygdala, and orbitofrontal cortex^7,8^. Behavioural and neuroimaging research suggests that the perception of visual, auditory and somatosensory inputs is affected by the heart phase^2,9,10^. These studies show that participants are more sensitive to exteroceptive sensory signals during diastole and less sensitive during systole when key sensory brain regions receive cardiac related afferent signals increasing the excitability levels in these regions.

Learning is affected by states of cognitive and physiological arousal that can fluctuate over time ^11^. Importantly, model estimates of signed and absolute PE can capture these fluctuations as learning progresses. Although model estimates are good predictors of behavioural changes ^12^, studies exploiting concurrent trial-by-trial physiological changes can offer additional explanatory power when analysing behaviours or neural data related to signed and absolute PE. For example, some studies have used changes in eye gaze or pupil dilation to disentangle attentional and learning processes involved in PE coding ^13^. Others have used single-trial variability in EEG to expose latent brain states related to absolute PE, thereby complementing more conventional model-based fMRI analyses. Using such trial-by-trial estimates has revealed the temporal and spatial neural correlates of these learning signals in human and animal brains ^14–17^. Signed PE-related activity has been reported in a number of brain areas but absolute PE-related activity has been most often linked to ACC and AI ^5–21^ -- in or adjacent to brain areas associated with cardiac-related activity ^23^. In addition, recent studies have shown that absolute PE-related activity appears shortly after the outcome while signed PE-related activity has a longer latency after the outcome ^24–26^.

Because absolute PE information is neurally encoded early after outcome onset and given the adjacency of brain areas encoding both saliency and cardiac activity, we hypothesised that cardiac signals might interact with the impact that the absolute PE, as opposed to the signed PE, has on learning. By analogy with the variation in the impact of near-threshold perception that occurs in relation to the cardiac cycle, we investigated variation in the impact of near-threshold absolute PEs on learning as a function of the cardiac cycle. We did this by capitalising on a whole-brain machine learning technique and high temporal resolution data; we exploit trial-by-trial variability in the cardiac related signal to investigate separately how changes in absolute PE and signed PE throughout the task are modulated (enhanced or decreased) by the two cardiac phases. We hypothesise that the timing within the cardiac cycle (e.g. whether a decision outcome occurs at cardiac diastole or cardiac systole) modulates the strength of the neural representation of the outcome. In line with previous evidence showing a better ability to detect information during diastole as opposed to systole ^27^, we hypothesise that near-threshold absolute PE events, regardless of their perceptual magnitude will be better represented at diastole than systole. If this is the case, then internal subjective representations of choice value will reflect time points within the cardiac cycle when they are constructed.

## Results

### Statistics of the reward environment predict learning

Participants carried out a reward-guided decision task inspired by previous credit assignment tasks ^28,29^. On each trial, subjects were shown two visual cues that are associated with different category-specific brain areas (a face and a house), and asked to predict which colour (orange or blue) was most likely to follow (Fig. 1A, B). Participants made choices by pressing corresponding left or right buttons to indicate their prediction of orange or blue. The actual outcome (a single colour) was then displayed. Participants were instructed that the chance of the correct colour being blue or orange depended only on the cue-outcome prediction strength and the recent outcome history. While participants performed the task, we recorded neural responses to heartbeats with EEG and ECG (Fig.1A) during the outcome which included a four-second period to ensure that multiple heart beats would be recorded (mean: 4.58, std+-0.85). To modulate learning throughout the task, participants performed tasks employing four association schemes presented in separate blocks. There were three predictive schemes with high associations between cues and colours (highly predictive anticorrelated, highly predictive correlated, and variable predictive schemes) and one scheme with no associations between cues and colours (non-predictive scheme) (Fig.1C).

**Figure 1.**
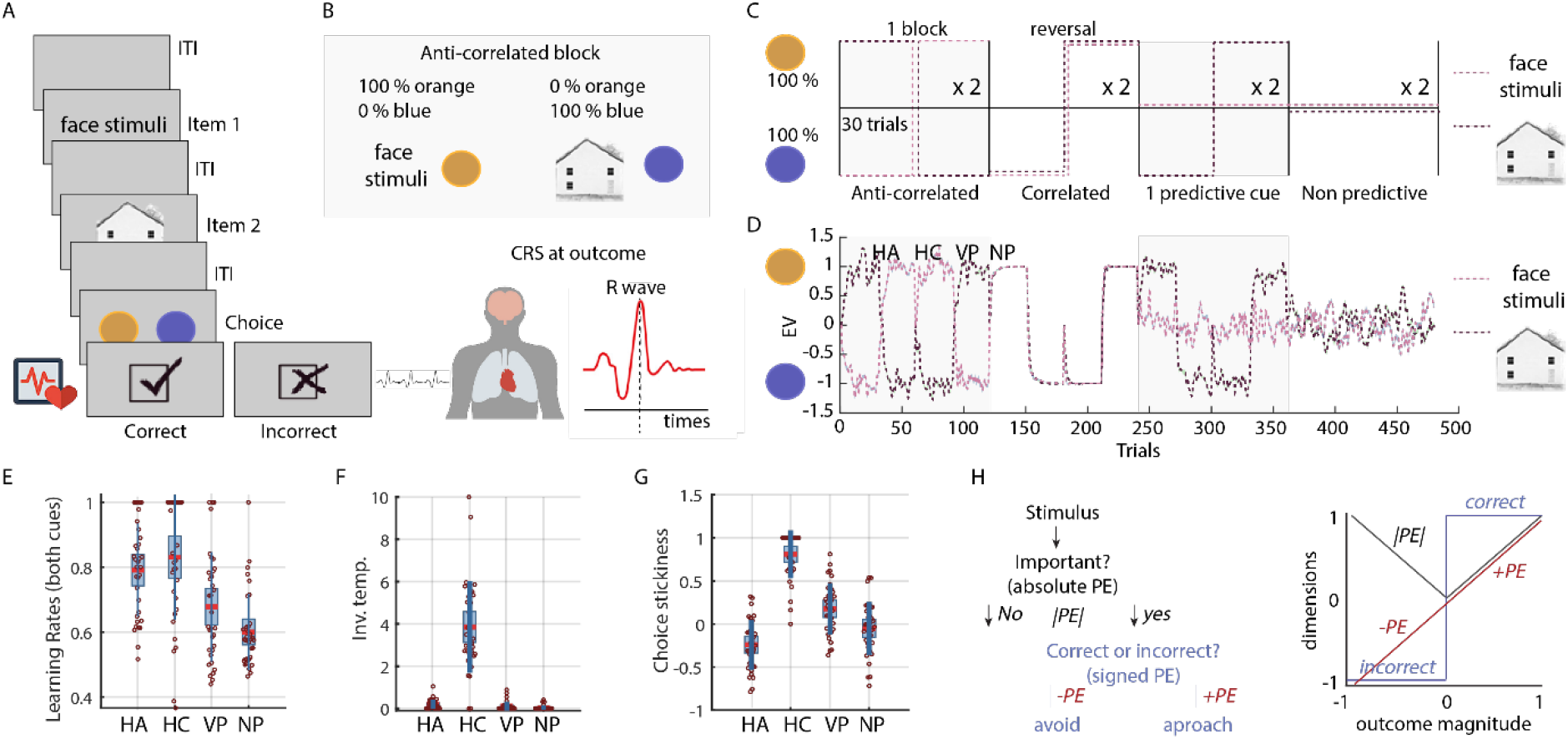
Schematic representation of the task and RL results for all four association schemes, highly predictive anticorrelated (HA), highly predictive correlated (HC), variable predictive (VP) and non-predictive (NP). **(a)** task + cardiac-related neural signals (CRS) recorded at outcome for 4 sec -enough to get on average 3.5 CRSs **(b)** example of association between cues and predicted colour for one version of the task (anti-correlated blocks – see Methods) **(c)** prediction strengths for each block **(d)** model prediction of choices **(e-g)** RL parameters **(h)** definition of absolute and signed PE

Initial analyses confirmed that the cue-colour prediction strength (which determined reward contingencies) was the primary modulator of adaptive behaviour in the task (Supplementary Figure 1). To investigate the two dimensions of learning (signed and absolute PE), we modelled participants’ choices using two canonical reinforcement learning models which differed in the way the cues’ identities were perceived (either a model incorporating biases for either faces over houses or *vice versa* and a model devoid of any such bias) resulting in different prediction weights associated with each cue (see Computational Modelling in Methods). A formal model comparison showed that the simpler prediction model, lacking any bias, outperformed the more complex version incorporating cue bias (BIC=362.1738, BIC=377.4045) and suggested that participants did not exhibit a bias towards one or other cue type. Having established the goodness of fit of the prediction model to behaviour, all further analyses were conducted using the outcome-related signals estimated with this model (Fig. 1H).

### Grand average modulation of cardiac-related neural signals in learning-related dimensions

We next looked for EEG signatures of the heartbeat evoked potential (HEP). Figure 2a presents the topographical characteristics of the HEP based on the averaged cardiac-related neural signals (CRSs) recorded during the outcome (see Methods for the construction of the CRS). A morphology analysis revealed that the HEP was widely distributed along fronto-central and centro-parietal areas including the following spatial regions (Fronto-central sites: F1, Fz, F2, FC1, FCz, CF2; Centro-parietal sites: C1, Cz, C2, CP1, CPz, CP2 as in Figure 2a). We then probed whether these EEG HEP signatures were modulated as a function of learning. To do this, we looked at two dimensions of learning as well as outcome valence. The two dimensions of learning included the fully parametric signed PE signal and the absolute PE also called salience (which connotes how surprising an outcome is). This approach has the advantage of looking at two orthogonal RL signals as seen in Figure 1H. The outcome valence was simply correct versus incorrect outcomes. The results of the cluster-based permutation analysis (see Methods) revealed an increased HEP amplitude for trials with negative signed PE in comparison to positive signed PE (Monte-Carlo *p* value = 0.004) between 198 and 252 milliseconds in the fronto-central sites (Figure 2b). The contrast between correct and incorrect trials revealed an HEP amplitude increase for correct trials (Monte-Carlo *p* value = 0.005) in the time range between 286 and 348 milliseconds in the centro-parietal sites (Figure 2c). When contrasting trials in the absolute PE domain, we found an increased HEP amplitude on the low surprising trials as opposed to the high surprising trials (Monte-Carlo *p* values < 0.003); these differences were observed in two clusters with latencies 222 - 252 milliseconds and 418 - 464 milliseconds, both in the centro-parietal sites (Figure 2d).

**Figure 2.**
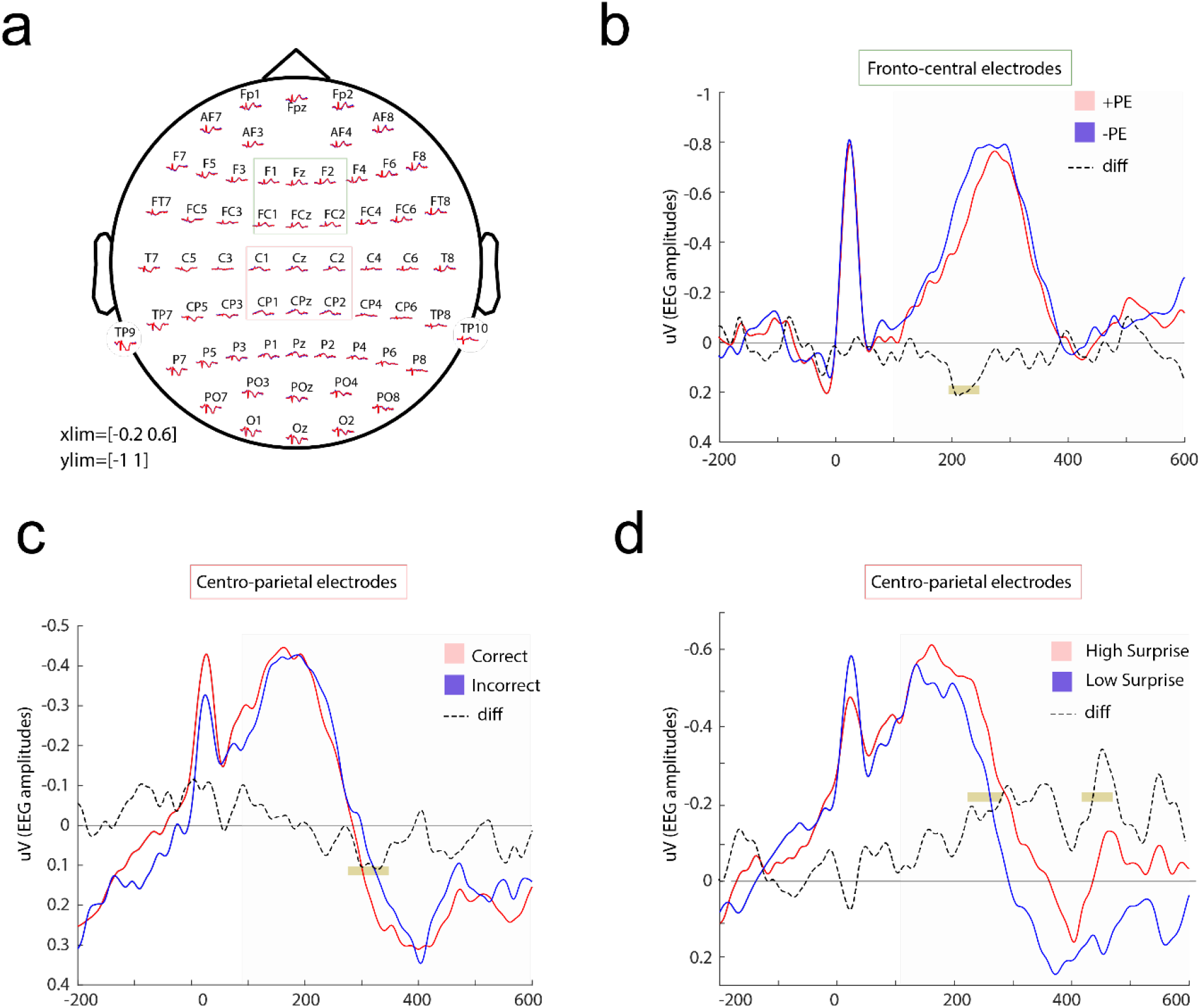
CRS morphology and results. **(a)** Grand average waveforms across the scalp time-locked to the onset of the R-wave which is the biggest electrical wave generated during normal conduction (time 0 ms, see Methods). The set of electrodes clustered by ROIs (colour-coded) for the fronto-central and central-parietal electrodes are represented for further analyses **(b-d)** CRS waveforms across all trials following the onset of the R-wave (at time 0 ms) are shown separately for **(b)** positive and negative signed PEs for the frontal cluster **(c)** correct and incorrect feedback for the central cluster **(d)** high and low surprising outcomes (absolute PEs) for the central cluster. The brown lines indicate the time-window (after the R-wave onset) for which a significant positive cluster was revealed by cluster-based permutation analysis. The dotted line represents the difference between the conditions (represented in red and blue). The shaded areas represent the time windows during which the analyses were performed and the HEP typically takes place (0.1-0.6s).

### The CRS is related to trial-by-trial variation in the absolute PE dimension and not signed PE

Rather than inspecting specific electrode averages, we next wanted to search for EEG heart-beat features that predict the learning axes. We thus moved on to identify the whole-brain heartbeat-evoked neural components of learning by using a multivariate single-trial discriminant analysis of the EEG (regularised Fisher Discriminant analysis, see Methods) of the CRS-locked signals. More specifically, for each participant, we used the average of the CRS for each outcome (see Fig.1a) and calculated the linear weights associated with each electrode that maximally separated (1) positive and negative signed PEs and (2) high versus low magnitude of absolute PE (i.e. the size of the unsigned PE which describes how surprising the outcome is). We did this over multiple temporal windows and quantified the classification performance by using the area under the curve (Az) using a leave-one out approach. This method has been well established in EEG data analysis ^30,31^. Using this machine learning approach, we showed the presence of a large CRS component reliably discriminating – even in individual participants (see Supplementary fig. 2) – between very high versus very low absolute PE outcomes. This component peaked in the time range 100–300 milliseconds after the R-wave (Fig. 3a-b). On the other hand, we did not observe any CRS component discriminating between positive and negative PEs (Fig.3f). By contrast, as noted, a grand average response difference was identified in the ERP in fronto-central electrodes (Fig.2b). In conjunction, the two results suggest that there is, on average, a difference in cardiac related signals when the valence of the signed PE is positive or negative^31^ but that trial-by-trial variability in the heart-related CRS does not reliably covary on a trial-by-trial basis with the trial-by-trial change in signed PE. In summary, the different analysis approaches (ERP and machine learning) suggest the possibility of a number of relationships between heard-related CRS and learning signals but converge in suggesting an especially clear link, even at the trial-by-trial level, between the heart-related CRS and absolute PEs connoting how surprising or salient an outcome is. We therefore focussed our analysis on the absolute PEs.

To test whether the absolute PE-related CRS was parametrically modulated by the model absolute PE (rather than responding categorically to very high vs. very low absolute PE), we then calculated the discriminator amplitudes for trials with intermediate absolute PE levels (i.e. low absolute PE [0.25 – 0.50]; and high absolute PE [0.5 – 0.75]) which were not originally used to train the classifier (also called the “unseen” data). To do so, we applied the spatial weights of the peak discrimination performance for the extreme outcome absolute PE levels to the EEG data with intermediate values. We expected that the discriminator amplitudes for these previously “unseen” trials would increase linearly as a function of absolute PE. Thus, the resulting mean amplitude at the time of peak discrimination would proceed from very low < low < medium < very high absolute PE. This is indeed what we found (Fig. 3c, blue: intermediate categories, grey: categories used for discrimination) confirming the linear relationship between the CRS-absolute PE component and its model-based counterpart (test on the left-out data: t31 =-7.3027; p=3.22e-8) and also the generalisability and robustness of our machine learning approach. Having applied the estimated electrode weights to single-trial data to produce a measurement of the discriminating component amplitudes (representing the distance of individual trials from the discriminating hyperplane), we thereafter used these amplitudes for all subsequent analyses involving single-trial CRS absolute PE.

Thus far, we have demonstrated that a large temporal component locked at R-wave onset within the CRS, the biggest wave generated during normal heart conduction (see Methods for full definition) discriminated all heart-evoked signals that were collected during the feedback period along the absolute PE dimension. However, the extent to which this temporal component was driven by the first, second, or third heartbeat that occurred in the outcome period remained unknown. Next, we therefore repeated our multivariate analysis independently for each of the three possible CRS times post-outcome, sequentially, across all trials to better understand the temporal dynamics of the absolute PE CRS modulation. This approach allowed us to determine which heartbeat was most related to absolute PE. Applying this method, we showed that only the first CRS after feedback contained information about absolute PE that could be revealed with machine learning techniques, in the range 100–300 milliseconds after heart beat onset (Fig. 3d). This finding indicates that the first heartbeat after outcome is the one that relates most to the representation of the absolute PE of the outcome, suggesting that the timing of the outcome with respect to the cardiac cycle might be important in determining how participants update their internal representations of decision outcomes.

### Effect of the cardiac cycle timing on the CRS absolute PE amplitude

Having identified a CRS component associated with absolute PE, we next asked whether the timing of the outcome along the cardiac cycle further modulated the amplitude of this signal. In other words, we examined whether the magnitude of the CRS representing absolute PE in the EEG epoch related to the first heartbeat after feedback onset would be higher or lower when the outcome was presented during diastole compared to systole. This would be an indication that internal subjective representations of how surprising an outcome is (compared to expectation), depends on the natural oscillation of the heart. To answer this question, we identified all outcomes with onsets which happened at diastole and all outcomes with onsets which happened at systole (see fig. 4a). This split allowed us to compute the mean CRS for heartbeats after outcomes presented at diastole, and after outcomes presented at systole. Importantly, although the outcome onset was defined according to the systole and diastole periods in the previous R wave, the absolute PE-related CRS components that we analysed were defined in the next R wave (see Methods). Naturally, as the diastole phase is longer on average than the systole phase, we expect a higher number of outcomes presented during the diastole phase (m=65±5 and 54±5 for diastole and systole, respectively). First, we confirmed that all other aspects of the task remain identical in these conditions. For example, the frequency of occurrence of both conditions was similar in the different learning blocks (e.g. predictive or non-predictive) employed in the task (see Methods; systole F_3_=0.2, p=0.893; diastole F_3_=0.2, p=0.896, supplementary Fig.3a) and were associated with similar levels of overall reward received in the task (t_31_=0.8046, p=0.4272, supplementary Fig.3b), unsigned PEs from the RL model (t_31_=-0.0581, p=0.9541, Supplementary Fig 3c) or signed PE (t_31_= -0.72, p= 0.4769).

Having split the outcomes according to the cardiac cycle, we found that the associated absolute PE-related CRS depended on whether the outcome was presented at systole or diastole, with the mean CRS magnitude related to absolute PE being more negative when outcomes were presented at the diastole compared to the systole phase (t_31_=2.8460, p=0.0078, fig4b). Additionally, beyond the linear relationship between absolute PE and CRS (fig3c), we found that the effect of the cardiac cycle increased as absolute PEs became smaller (main effect of absolute PE: t_252_=14.055, p=1.6e-33; heart cycle: t_252_=-2.154, p=0.0321; see fig4e). This effect was mainly driven by the fact that near-threshold absolute PE were more strongly represented at diastole (t_31_=2.4089, p=0.0221, fig4c). This confirms that timing within the cardiac cycle modulates neural signals of absolute PE and that these representations are stronger after an outcome is presented at diastole compared to systole.

We then asked whether these differences in the way naturally occurring bodily oscillations modulate the neural signals that determine learning can also impact participants’ decisions (which should be guided by learning that is based on the same neural signals). As we had found a more negative CRS when outcomes were presented at diastole compared to systole, we wondered whether inter-individual differences in the impact of the cardiac cycle on the CRS component co-varied with task performance and learning. We thus first ran a regression analysis for each participant to test the extent to which the cardiac cycle influenced their neural activity. Participants with a higher regression coefficient would have a stronger decrease in the CRS for outcomes presented at diastole compared to systole (see figures 4f-k). We expect these participants to be the ones showing a greater propensity for learning as their sensitivity to near-threshold events would be enhanced. As a consequence, they should also be the ones who ultimately receive more rewards overall. To test this hypothesis, in a second step, we ran a correlation between the regression coefficient and the mean reward and learning rates in the task. In line with our predictions, we found that participants showing a higher difference in CRS between diastole and systole were the participants that had higher learning rates and better task performance as indexed by the total number of rewards received (learning rates: t30=2.1821, p=0.0371; reward: t_30_=2.4329, p=0.0212; Fig.4f and 4g). In summary, we can link interindividual variation in cardiac modulation of learning signals to interindividual variation in the parameters of a computational model of learning and to individual variation in an index of behaviour – overall rewards – independent of the computational model. To further examine the relationship between diastole-based CRS-related neural indices and learning, we examined task blocks where learning was possible (predictive blocks) or not possible (non-predictive blocks). The relationship was only present in the blocks in which learning was possible (*predictive blocks:* learning rates: t_30_=2.4468, p=0.0205; reward: t_30_=2.3751, p=0.0241 Fig.4g and 4j; *non-predictive blocks:* learning rates: t_30_=-0.4445, p=0.6598; reward: t_30_=1.1354, p=0.2652 Fig.4h and 4k). These results remain true even when including a covariate indexing features of the external outcome type – reward and absolute PE from the model as opposed to the internal, subjective, absolute prediction error (see supplementary figure 4).

## Discussion

In this new study, we combined machine learning-based analysis techniques and EEG to investigate the contribution of cardiac related neural signals on several dimensions of reward-based learning. We first show the intrinsic relationship between the absolute PE dimension of decision outcome and the CRS. In addition, we demonstrate that the timing of a reward-related outcome, with respect to the cardiac cycle, shapes the CRS magnitude as well as learning and overall performance in the task. More specifically, absolute PE discrimination computed during the first CRS after outcome, was lower if the outcome happened at systole than diastole and this difference, across participants. This difference was related to learning rates in the computational model and ultimately performance as indexed by a simple, computational model independent measure – the number of rewards received – in the task. Furthermore, the relationship between single-trial CRS and learning was only observed in the block of the task where learning was possible.

Our results demonstrate that single-trial CRS recorded during the presentation of reward-related outcomes discriminates between different levels of absolute PE outcome. By contrast, the magnitude of the CRS did not differ when contrasting positive *versus* negative signed PEs. The absolute PE and signed PE components of reward learning subserve different functional roles in learning ^5,24^; whilst signed PE is associated with approach-avoidance behaviour, absolute PE, also called salience impacts future attentional engagement; an effect that is determined by the magnitude of the discrepancy between prior expectations and outcome. Cardiac neurophysiological responses often convey not only information about the current bodily state, but they also carry predictions of how the bodily system should organise internal resources to deal with expected future sensory information ^27,32^. These cardiac predictions are often accompanied by a modulation of attentional responses to upcoming stimuli that, ultimately, are homeostatically relevant. In this way, it has been suggested that the internal bodily state determines perceptual stimulus salience in relation to homeostatic levels ^32–34^. For example, a stimulus occurring when resources are sparser may be perceived as more salient than a stimulus occurring when more resources are available. Here, we found a strong relationship between outcome absolute PE and the CRS magnitude, suggesting that the resulting CRS-related component might signal how much attention needs to be deployed to the current outcome. In this way a bodily signal might modulate learning.

Neuronal models of interoception conceptualise cardiac predictions as afferent signals projecting between agranular visceromotor areas in frontal and prefrontal cortices and the anterior insula, which serves as the primary interoceptive cortex ^35,36^. The anterior insula is argued to be a main neural source for the CRS along with other interconnected areas such as the cingulate and the somatosensory cortices ^37–39^. These brain regions belong to a wider network, often referred to as the salience network, which is sensitive to homeostatically relevant stimuli independent of whether their valence is negative (penalising) or positive (reinforcing)^18^. It is becoming increasingly clear that neural responses in the absolute PE network rise quickly after an outcome is revealed ^24–26^. Here we observe that the CRS is parametrically modulated by the outcome’s absolute PE and that this is mainly due to the first heartbeat recorded immediately after the outcome onset. This means that CRS magnitude changes recorded immediately after outcome can be used as a proxy for attentional allocation of the internal representation of absolute PE, highlighting the unique and crucial contribution of cardiac brain responses in reward learning.

Previous studies have carefully time-locked the presentation of stimuli to the cardiac phase to investigate differences in the way they are processed ^2,40,41^. However, sensory or learning information is not presented in such a phase-locked manner in our everyday lives. By investigating how participants naturally receive information relevant for learning and assign credit for outcomes to objects maintained in memory, with respect to the natural timing of the cardiac cycle, we have adopted an ecological approach to studying brain-heart interactions in the context of learning and decision making. Previous studies, adopting a similar approach, have shown that people actively seek information in the world, or more precisely sample the world through active sensing, as a function of the cardiac cycle. For example, in an active sampling visual paradigm, saccades and visual fixations are more likely to occur in the quiescent phase of the cardiac cycle (e.g. diastole) ^42^. Similar work suggests that people actively adjust sensory sampling so that more time is spent in the diastole period in which perceptual sensory sensitivity is enhanced ^43^. In our study, we have shown that the magnitude of the CRS is stronger when the outcome appeared during the diastole period in comparison to the systole period (Fig.4). This suggests that the phase of the cardiac cycle is an important modulator of internal representation and cognition and influences the way in which we naturally receive information.

Importantly, we also observed that the influence of the cardiac cycle on the absolute PE CRS magnitude progressively increased as the outcome absolute PE became smaller. In outcomes with near-threshold absolute PEs, the CRS magnitude increase was predominantly observed when the outcome was presented at diastole (Fig.4). This means that when the decision maker’s prior expectations are close to the outcome (i.e. small adjustments between expectations and outcomes) learning is more likely to occur during the quiescent phase of the cardiac cycle than during the active, systolic phase. Neuronal excitability is influenced by the cardiac cycle; whilst neural signals from the baroreceptors occurring at systole attenuate concurrent brain activity ^12, 13^ and impair information processing, enhanced excitability and perceptual processing is observed during diastole ^14, 15^. Formally, enhanced neuronal excitability may increase neural gain, which directly translates into an increase of the breadth of attention towards the aspects of the environment to which one is predisposed to attend ^16^. Here we show that in instances where learning happens in small increments because the PE-related surprise is not very salient, learning is enhanced during diastole compared to systole, helping to update prior expectations even when there is little new information available.

Beyond showing modulations of the CRS amplitude timed to different phases of the cardiac cycle, our results demonstrate that these heart cycle-specific neuronal changes translate into individual differences in overall learning. Individuals that exhibited higher differences in the absolute PE CRS magnitude changes to outcomes presented at diastole versus systole also showed higher learning rates and better overall task performance. Individual differences in cardiac neural responses have long been established ^17^. For example, HEP amplitude modulation often present during observation of highly salient stimuli is stronger for individuals with greater self-reported empathy scores ^18^. Also, individuals with low cardiac interoceptive sensitivity show greater difficulty retrieving information presented at systole in comparison to those with high interoceptive sensitivity ^19^. Additionally, we found that these individual differences in the effect of the cardiac cycle on absolute PE encoding were only true in task blocks where learning was taking place versus blocks where learning was precluded (i.e. random contingency between colours and stimuli). Increased and decreased cardiac sensitivity has also been shown to help or hinder adaptive intuitive decision making when the generated cardiac predictions favour advantageous choices - i.e., when learning is taking place; however, the opposite is true when predictions are towards disadvantageous choices ^20^.

Our finding that absolute PE representation depends on the heart cycle might also be described in terms of periodical modulations of internal value representations in a predictive coding framework. According to this framework, the brain is constantly creating and updating predictive internal models of sensory inputs, including both exteroceptive and interoceptive signals such as the heartbeat. As each heartbeat and its accompanying pulse wave cause temporary physiological changes throughout the body, the brain treats these recurring cardiac signals as predictable events and attenuates them to reduce the chances of mistaking these self-generated signals for external stimuli ^44–46^. As a consequence, for example, in the context of somatosensory events, sensory discrimination is less accurate during systole than diastole ^43^. However here for the first time, we have shown that even if the sensory information is the same, the extent to which an absolute PE affects learning is also linked to the cardiac cycle; the degree to which an internal model of a cue-outcome association is strengthened depends on the cardiac cycle, in line with a predictive coding account for cardiac phase-related internal fluctuations.

## Methods

### Participants

Thirty-five healthy, right-handed adults participated in the experiment. 3 participants were excluded due to excessive noise in the EEG signal so that data from 32 participants were included in the analyses (24 ± 7.13; 10; 0.83 ± 0.13; where numbers correspond to mean age ± SD; number of female participants, handiness mean ± SD; as measured by the Edinburgh handedness inventory ^21^). All participants were naïve to the task, had no personal or familial history of neurological or psychiatric disease, were right-handed, gave written informed consent (Medical Science Interdivisional Research Ethics Committee, Oxford RECC, No. R55856/RE002), and received monetary compensation for their participation. Sample sizes were determined based on previous studies that have used similar reward learning paradigms to investigate brain responses during learning ^28,30,47^, and studies that have measured the CRS to investigate neural responses to heartbeats in humans ^14, 22^.

### Stimuli

Stimuli consisted of pictures of 10 faces and 10 houses (512 × 512 pixels) adapted from previous EEG experiments^48^, two circles in blue and orange (125 × 125 pixels). All the stimuli were equalised for luminance and contrast. The outcome images consisted of a tick and a cross, which were also equalised for luminance and contrast.

### Experimental design

Participants were seated in a dimly lit, sound-attenuated, and electrically shielded chamber in front of a monitor at a distance of 70 cm. EEG was recorded using a 64 channels cap (see EEG data collection section) while participants performed a reward-based learning task. Participants’ ECG was recorded with a standard EEG electrode attached to their chest to monitor heart activity throughout the session. The experiment consisted of eight blocks of 60 trials (480 trials in total) separated by small breaks. At the beginning of each block, the association between colours and objects changed. Two new objects were presented in each block: a house and a face. Each object was uniquely associated with a colour according to different schemes. There were a total of four types of blocks: three predictive blocks (in which objects predicted outcomes) and one non-predictive block (from which predictive associations between objects and outcomes were absent). The three predictive blocks contained the following associations: (1) both stimuli were highly predictive and there was a negative correlation between each stimulus and respective associated outcomes (i.e. each stimulus predicted a different colour), (2) both stimuli were highly predictive and positively correlated in the outcomes that they predicted (i.e. both stimuli predicted the same colour), (3) only one stimulus was highly predictive and the other non-predictive. In the non-predictive block, the two objects were not associated with any colours.

### Learning task

We used a modified version of the weather prediction task. In a typical version of the task, participants have to predict the weather (rain/sun) on the basis of probabilistic cues. To avoid any subjective preference, we changed the sun/rain to two neutral colours (light blue/orange). We also presented one object at a time to isolate the EEG responses to faces and houses. On each trial, participants first saw a fixation cross for 500ms, followed by the presentation of one stimulus that could be either a face or a house (500ms). This was repeated for the second object (same timing). Each possible pair of objects: Face-House, House-Face, Face-Face and House-House were presented equally often and counterbalanced across a block. After the presentation of both objects, participants had to make a decision between two colours, orange and blue on the basis of their estimates of the association between the objects and colours as well as on the basis of what the particular combination of objects would be likely to predict. For example, if the house predicted orange deterministically (100%), the face predicted blue and they were presented together, then there was a 50/50% chance of getting a blue/orange. If, however, the face was presented twice, then the outcome was blue, 100% of the time. The decision phase lasted 1200ms. After participants made their decisions, they saw the outcome of their choice for 4000ms, which allowed us to record on average four heartbeat per outcome (mean: 4.58, std+-0.85 across trials). After the task, participants were given a debrief and paid £20 for their participation. They were told that they would receive a fixed payment for participation (£15 per hour) and an additional amount (up to a maximum of £5) based on the outcome of a random subset of trials selected at the end of the experiment (excluding ‘lost’ trials). No further details regarding the mapping between earned points and the final payoff were given to the subjects.

### Computational modelling

#### First model

We used a canonical Rescorla-Wagner Model that computed a Prediction Variable as follows: *PV*_*n*_ *= 0*.*5 x V1*_*n*_ *+ 0*.*5 x V2*_*n*_ where *V1*_*n+1*_ *= V1*_*n*_ *+* α *x PE*_*n*_ and *V2*_*n+1*_ *= V2*_*n*_ *+* α *x PE*_*n*_ where PV sums up the equally weighted stimulus–outcome association strengths for each item (faces or houses; V1 and V2 that could either be Face-Face or Face-House or House-House). PE is the prediction error. PV is then converted to a choice probability following the equation: *p* = 1/(1 + *e*(β * (PV – 0.5) + γ * *Cn*−1)), where β is the inverse temperature, or exploration parameter and γ represents the choice stickiness^49,50^ (the degree to which choices are likely to simply be repeated from trial to trial regardless of outcome). *Cn*−1 is the choice in the previous trial (orange choice coded as +1 and blue choice coded as −1). *V* is the item–outcome association strength of each item, *O* is the outcome in the current trial (orange outcome coded as +1 and blue outcome coded as 0), and α is the learning rate shared by both items. The subscript *n* represents the current trial, and *n* + 1 represents the updated trial. This model is based on the empirically validated assumption that the prediction used at time of decision is calculated at the outcome phase ^28^. There are three free parameters in this model: the learning rate α, the exploration parameter β, and the choice stickiness factor γ.

#### Second model

The second model resembled the first but also allowed a different weight for the two stimuli when the prediction variable is being calculated at the time of decision. This was achieved by inclusion of an additional free parameter called the prediction weight (*pw*). The weight associated with each cue is either the weight for the stimulus selected or the non-selected one. This model makes it possible to capture any bias the participant might have to rely more on predictive association relating to either the face or house stimulus. We refer to this model as the Prediction Weight Model (Model 2).

#### Model comparison

We formally compared the two models by examining how well they fit the behavioural data collected. We utilised a maximum likelihood estimation and a constrained non-linear optimization approach (as implemented in *fmincon* in MATLAB R2020) individually for each session to estimate the three free parameters (α, β, γ). It is important to note that there was no constraint on the fitted parameters from the other participants or the group of other participants so that the fitting procedures for each individual’s data were independent of each other. In addition, the estimated values of fitted parameters and negative log likelihoods were stable across the fitting procedure given different initial values. The negative log likelihoods estimated for individual participants were transformed to calculate Bayesian Information Criterion (BIC) values by including the number of free parameters. The subsequent comparison was based on the sum of these two estimated BIC measures, which penalised the inclusion of additional free parameters.

### EEG and ECG recording

EEG was recorded with sintered Ag/AgCl electrodes from 62 scalp electrodes mounted equidistantly on an elastic electrode cap (64Ch-Standard-BrainCap for TMS with Multitrodes; EasyCap; two cap sizes, 56 cm and 58 cm head circumference). The distance between electrodes was on average 3.3cm and 3.5cm for the 56cm and the 58cm cap, respectively. The Ground electrode was located centrally at the electrode site corresponding to AFz in the 10/20 system. An additional ECG electrode was placed on the participants’ chests around 12 cm below the left clavicle. All electrodes were referenced to the right mastoid and re-referenced to the arithmetic average reference of all electrodes off-line. Continuous EEG was recorded using BrainAmp amplifiers (BrainProducts, Munich, Germany; 0.1 μV analog-to-digital conversion resolution; 1000 Hz sampling rate; 0.01-100 Hz online cut-off filters).

### EEG data analysis

Off-line EEG analysis was performed using Fieldtrip (https://www.fieldtriptoolbox.org/). The data was digitally band-pass-filtered between 0.5-40 Hz. Bad/missing channels were restored using a FieldTrip based spline interpolation (1-2 electrodes per participant on average). Detection of R-peaks in the ECG recording was done using the Pan-Tompkins algorithm as implemented in MATLAB ^23^. Next, the data were segmented into intervals time-locked to either the onset of the feedback, or the R-peak onset of the heartbeat R-waves occurring during the feedback period, or the onset of the visual images (faces/houses). The R-wave is the biggest wave (indicating the changing direction of the electrical stimulus as it passes through the heart’s conduction system) generated during normal conduction and the first upward deflection after the P wave part of the QRS complex as presented in figure 4a. The R-peak of the R-wave determines the time 0ms of our CRS.

The intervals time-locked to the feedback onset were segmented into 4.9s intervals starting from 0.9s before the feedback onset. The intervals time-locked to the onset of the R-wave were segmented into 0.8s intervals starting from 0.2s before the R-wave onset. The intervals time-locked to the onset of the visual images were segmented into 0.7s intervals starting from 0.2s before the stimulus onset. This was done separately for positive versus negative PEs, high versus low absolute PE, and for correct versus incorrect trials.

Automatic artefact rejection was performed excluding trials and channels whose variance (z scores) across the experimental session exceeded a threshold of 20 μV. This was combined with visual inspection for all participants eliminating large technical and movement related artefacts. Physiological artefacts such as eye blinks, saccades and the volume-conducted cardiac-field artifact (CFA) were corrected by means of independent component analysis (RUNICA, logistic Infomax algorithm) as implemented in the FieldTrip toolbox. Those independent components (4.78 on average across participants; 1.13 SD) whose timing and topography resembled the characteristics of the physiological artefacts were removed. The CFA represents a challenge to the analysis of the HEP because the averaging of the data around the R-peak amplifies the CFA becoming time-locked to the heartbeat ^51^. Nonetheless, ICA has been shown to be of high efficiency in the removal of the independent components representing CFAs from the EEG signal ^37,52,53^. The IC identification and selection process were guided by visual inspection of their properties, based on time course and scalp topography. ECG channels were excluded from the analysis and the signal was then re-referenced to the arithmetic average of all electrodes.

For the ERP analysis, the segments were baseline-corrected using an interval from -0.15s to -0.05s for the segments time-locked to the R-wave onset, an interval from -0.9s to -0.1s for the segments time-locked to feedback, and interval from -0.2s to -0.05s for the segments time-locked to the visual stimulus onset. To further ensure that the HEP changes that we observe are not influenced by CFA artifacts, and they are truly locked to the participants’ heartbeat, we created surrogate R-peaks by shifting the onset of the original R-peak ^8, 9^. R-peaks were shifted within a time window of −500 to +500 ms and they were shifted by the same amount separately for each subject and each of the four learning blocks. We subsequently applied the same criteria for calculating HEP amplitude and submitted these surrogate values to the cluster-based permutation test as described below.

### Topography and statistical analysis of the ERPs

In light of the considerable variability in the polarity, latency and scalp distribution of the HEP [9, 10] we adopted a non-parametric, cluster-based permutation approach to first determine the HEP morphology, and then estimate any HEP amplitude modulation as a function of learning. Subject-wise activation time-courses were extracted and passed to the statistical analysis procedure in FieldTrip, the details of which are described by ^54^; Subject-wise activation time-courses were compared to identify statistically significant clusters in the time and spatial domain using a FieldTrip-based analysis across all time points and electrode sites. FieldTrip uses a nonparametric method ^55^ to address the multiple comparison problem. T-values of adjacent temporal and frequency points whose p-values were less than 0.05 were clustered by adding their t-values, and this cumulative statistic is used for inferential statistics at the cluster level. This procedure, i.e. the calculation of t-values at each temporal point followed by clustering of adjacent t-values, was repeated 5000 times, with randomised swapping and resampling of the subject-wise time-frequency activity before each repetition. This Monte Carlo method results in a nonparametric estimate of the P-value representing the statistical significance of the identified cluster.

The topographical distribution of the neural phenomena comprising the HEP was defined by computing mean voltages of the HEP time-locked to R-wave onset for all trials at the group level using the cluster-based permutation test including all electrodes sites and across the entire time window where the HEP typically takes place, this is, 0.1-0.5s ^22 2, 24 25^. In this analysis, no a-priori electrode clusters were formed (all active electrodes were treated as a distinct variable). The topography analysis revealed a number of electrodes widely spread along the frontal, centro-frontal and posterior areas where the HEP was distributed. These electrodes were then organised in 2 ROIs, a fronto-central ROI and a centro-parietal ROI, according to their spatial distribution (Figure 2.A) for further processing.

Next, we used the cluster-based permutation approach as implemented in Fieldtrip (see below) to test if HEP varied across the two main dimensions of learning: signed and absolute PE as well as correct versus incorrect outcomes. Since this method allows the comparison of only two conditions, we first organised the trials in two categories. We thus computed averaged signals aggregating trials with positive PE *versus* negative PE; and trials with high absolute PE *versus* low absolute PE and trials with correct *versus* incorrect outcome. Thereafter, we ran three parallel contrasts on averaged HEP contrasting trials with correct *versus* incorrect valence; trials with high positive PE *versus* negative PE; and trials with high absolute PE *versus* low absolute PE, by means of within subject non-parametric cluster-based permutation analysis as described above and represented in Fig.2b. A non-parametric, cluster-based permutation approach is an efficient way of dealing with the multiple comparison problem that prevents biases in pre-selecting time-windows avoiding inflation of type I error rate. Thus, the statistical analyses were performed across the entire time window in which the HEP typically takes place (0.1-0.6s) and restricted to the ROIs defined according to the HEP morphology analyses. For each comparison, subject-wise activations at electrode sites circumscribed in the ROI were extracted and passed to the analysis procedure. To avoid spurious findings, significant effects of 15 milliseconds or shorter were discarded from further analysis. Where appropriate, p-values were corrected for multiple comparisons using Bonferroni-Holms correction.

### Multivariate analyses

We hypothesised that the CRS, that is, the epoched EEG data synchronised to the heart at time of outcomes, may be associated with reward-based learning dimensions. To investigate this idea, we used a linear multivariate classifier, with a sliding window approach, on the CRS data. Specifically, we searched for a projection of the multidimensional EEG signal, xi(t), where i={1…T} and T is the total number of trials, within short time windows that achieved maximal discrimination between binary groups of trials as described in ^26,31^. Locked to the heartbeat, these binary groups included: (1) positive versus negative signed PEs, (2) very high and very low absolute PEs.

All analyses were performed on windows with a length of N=60 ms and the window centre τ was shifted from −100 to 600 ms relative to the heartbeat onset, in 10-ms increments. We applied a regularised Fisher discriminant analysis to find the spatial weighting, w(τ), that maximally discriminated between the binary groups described above, arriving at a one-dimensional projection yi(τ), for each trial i and a given window τ:

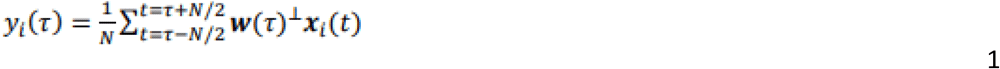

where yi(τ), is organised as a vector of single-trial discriminator amplitudes (1 × Trials), the spatial filter, w(τ), is organised as a vector with as many weights as there are channels in the data (1 × 64) and data, xi(τ), is organised as a matrix, with dimensions (64 × Trials/Samples). We adopted this approach to identify all time windows τ yielding significant discrimination performance in the heart-related period. The projection vectors w at each time window τ were estimated as: w=Sc(m_2_–m_1_) where *m*_*i*_ is the estimated mean of condition i and Sc=1/2(S_1_+S_2_) is the estimated common covariance matrix (that is, the average of the condition-wise empirical covariance matrices, with T=number of trials). To treat potential estimation errors, we replaced the condition-wise covariance matrices with regularised versions of these matrices, with λ∈[0, 1] being the regularisation term and νthe average eigenvalue of the original Si (that is, trace(Si)/62). Note that λ=0 yields unregularized estimation and λ=1 assumes spherical covariance matrices. Here we optimised λ for each participant using a leave-one-out trial cross validation procedure across the entire post-outcome period.

We quantified the performance of the discriminator for each time window using the area under a receiver operating characteristic curve, referred to as an Az value, using a leave-one-out trial procedure. To assess the significance of the discriminator, we used a bootstrapping technique where we performed the leave-one-out test after randomising the trial labels. We repeated this randomization procedure 1,000 times to produce a probability distribution for Az, and estimated the Az leading to a significance level of P<0.01.

Given the linearity of our model, we also computed scalp topographies of the discriminating components resulting from equation (1) by estimating a forward model as:

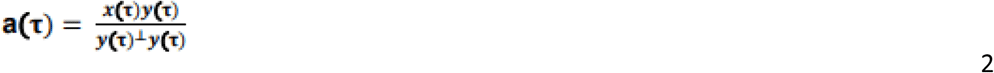

where yi(τ) is now shown as a vector y(τ), where each row is from trial i, and xi(τ) is organised as a matrix, x(τ), where rows are channels and columns are trials, all for time window τ. These forward models can be viewed as scalp plots and interpreted as the coupling between the discriminating components and the observed EEG.

### Diastole versus systole definition

Considering the biphasic nature of cardiac activity, we compared the cardiac neural response to absolute PE between the systolic and diastolic ventricular phases, namely, for simplicity, systole and diastole. We defined systole as the time between the R-peak and 300ms after R-peak (to coincide with the end of T-wave) (figure 3a) ^7,56^. We used the systolic offset of each cardiac cycle to define the onset of the diastole period, which ended at the R-peak. The non-equal length of systole and diastole meant that we were more likely (∼60%) to have an outcome onset in the diastole phases of the cardiac cycle. Each outcome was categorised depending on whether the stimulus occurred during systole or diastole. The average number of trials categorised as systole was 54.57 and as diastole was 65.35 with standard deviation of 4.99 and 5.05 respectively. Importantly, when an outcome was assigned to systole or diastole, the assignment depended on that outcome’s timing with respect to a current R wave. However, the absolute PE related CRS that were used for analysis in this work (figure 4) related to the next R wave.

**Figure 3.**
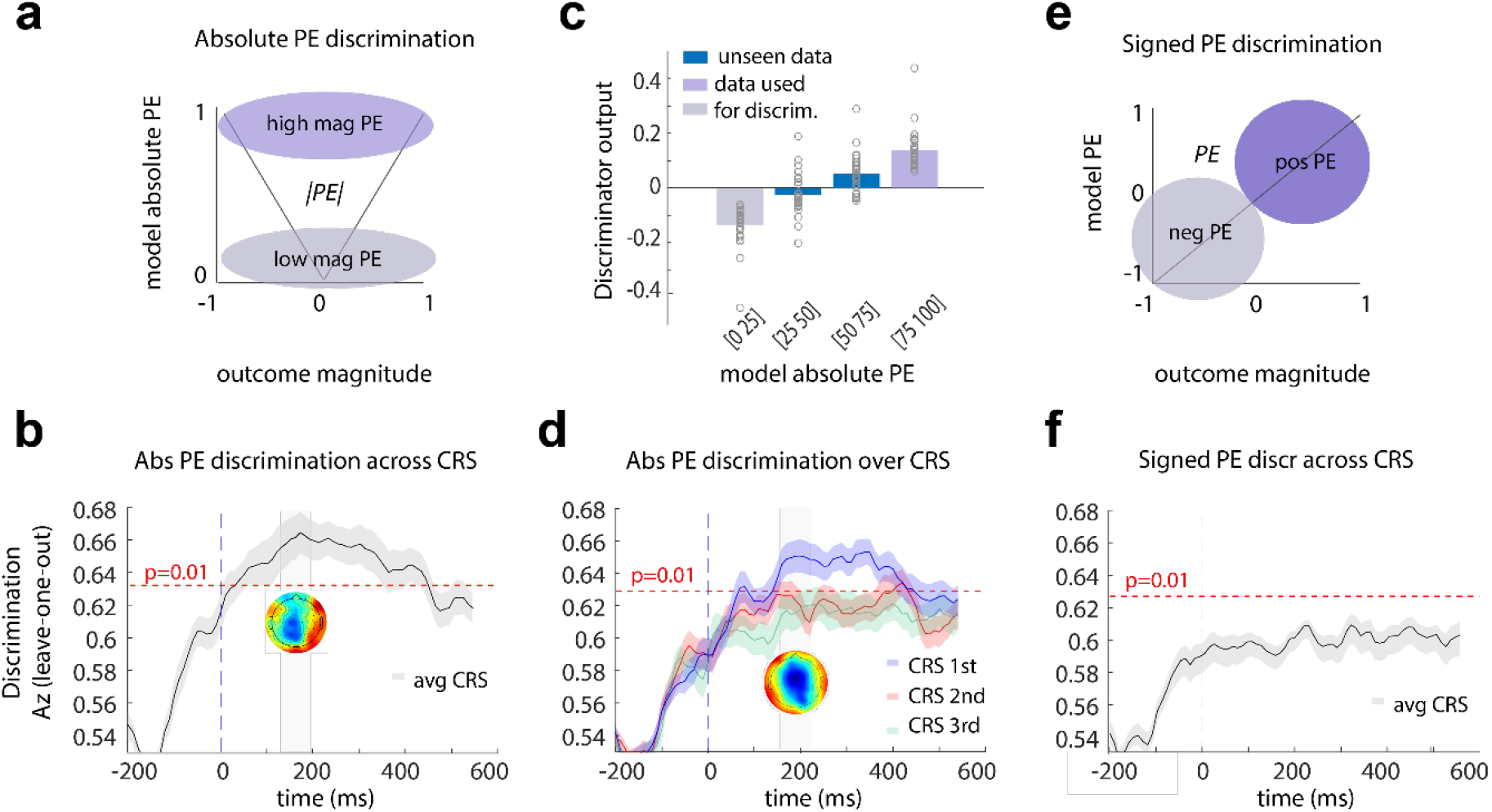
Machine learning discrimination. **(a)** Description of the data used for the outcome absolute PE discrimination: we used the highest and lowest quantiles based on absolute PE (salience or surprise) as estimates for high and low absolute PE respectively. The analysis was performed on the CRS-locked EEG data. **(b)** Discriminator performance (cross-validated Area under the curve, Az) during absolute PE discrimination (low vs. high surprising outcomes), averaged across subjects (N = 32). The dotted line represents the average Az value leading to a significance level of P = 0.01, estimated using a bootstrap test. Shaded error bars are standard errors across subjects. This analysis used all first three CRSs after outcomes. The scalp map represents the spatial topography of the absolute PE component. **(c)** Mean discriminator output (y) for the absolute PE component, binned in four quantiles based on model-based absolute PE estimates, showing a parametric response along the absolute PE dimension. Purple bins indicate trials used to train the classifier, while blue bins contain “unseen” data with intermediate absolute PE levels. Points are individual subjects. **(d)** The same analysis as in figure 3b (N = 32) for each CRS after outcome. Blue represents the first heartbeat after outcome, red the second and green the third. The dotted line represents the average Az value leading to a significance level of P = 0.01, estimated using a bootstrap test. Shaded error bars are standard errors across subjects. The scalp map represents the spatial topography of the absolute PE component. **(e)** Description of the data used for the signed PE discrimination: we used outcomes defined by the RL model as either positive or negative PEs. **(f)** Discriminator performance (cross-validated Az) during signed PE discrimination (positive versus negative PE), across all CRS after outcome, averaged across subjects (N = 32). The dotted line represents the average Az value leading to a significance level of P = 0.01, estimated using a bootstrap test. Shaded error bars are standard errors across subjects. No components were identified.

**Figure 4.**
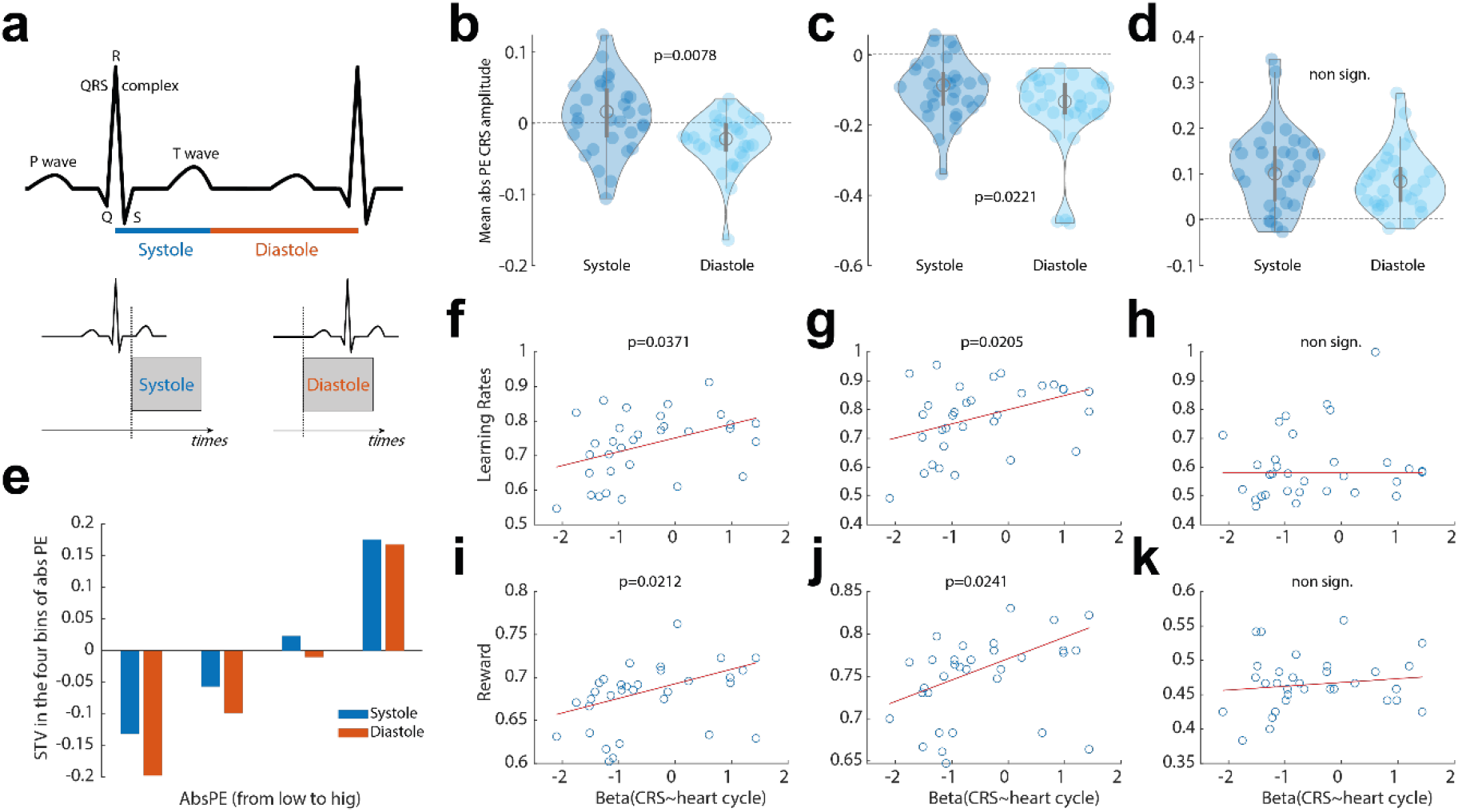
Influence of the cardiac cycle on the CRS and learning. **(a)** Schematic description of the systole and diastole phases. In red and blue are the systole and diastole periods respectively. Below is a representation of two example trials on which outcome onsets happened at systole or diastole. We then looked at the EEG response locked to the next heart-beat. **(b)** Mean amplitude difference between the CRS of the first heartbeat after all outcomes presented at systole versus diastole (N=32). A violin plot is used to present all individual participants’ averages. **(c)** Mean amplitude difference in the CRS for the low salient outcomes presented at systole versus diastole across subjects (N=32). **(d)** Mean amplitude difference in CRS for the high salient outcomes presented at systole versus diastole (N=32). **(e)** Parametric effect of the CRS showing that the lower the absolute PE, the more there is a significant difference between the CRS at diastole and systole. **(f)** learning rates – all task blocks **(g)** learning rates - predictive blocks **(h)** learning rates - non predictive blocks **(i)** reward – all task blocks **(j)** reward - predictive blocks **(k)** reward - non predictive blocks. **(f-k)** In red, the fit of the robust regression. Any of these results remain true even when including a covariate indexing features of the external outcome type – reward and absolute PE from the model.

### Regression analysis

To examine the association between the cardiac cycle (i.e. diastole: 1, and systole: 0) and the neural cardiac-related signal, we performed the following logistic regression analysis (separately for each participant):

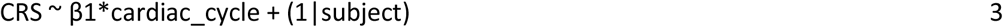

We then tested whether the regression coefficients across participants (β1 values in Eq. 3) came from a distribution with a mean different from zero (using a t test). To control for potential confound of outcome, we also performed the following logistic regression analysis (separately for each participant):

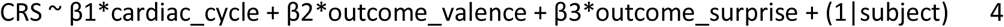

We then tested whether the regression coefficients across participants (β1 values in Eq. 4) came from a distribution with a mean different from zero (using a t test).

## Acknowledgement

Funding for this work was provided by the UKRI to Elsa Fouragnan (MR/T023007/1), the Bial Foundation to Alejandra Sel (Grant 44/16) and the Wellcome Trust to Matthew F. Rushworth (WT100973AIA). We also thank Miriam Klein-Flugge for helpful comments on the manuscript.

## Author’s contribution

E.F., M.R, A.S. designed the experiment; Y.C., B.P., A.S. collected the data; E.F., B.H, A.S. analysed the data; E.F., M.R. and A.S. wrote the manuscript. All authors discussed the results and implications and commented on the manuscript at all stages.

## Conflict of Interest Statement

The authors declare that the research was conducted in the absence of any commercial or financial relationships that could be construed as a potential conflict of interest.

## Data availability

We have deposited all choice raw data in an OSF repository. All reinforcement learning results in this paper are derived from these data alone. We have also deposited all neural data presented in the manuscript in the same OSF repository. The accession code is: https://https://osf.io/qgw7h/

## Code availability

The above repository also comprises the full reinforcement-modelling pipeline including model comparisons implemented in Matlab. All variables used for the EEG analyses are derived from this pipeline. Accession code to the repository is the following and a README inside the repository explains the details of its use: https://github.com/efouragnan/EEG-CRS_learning

## Supplementary figures

**Supplementary Figure 1.**
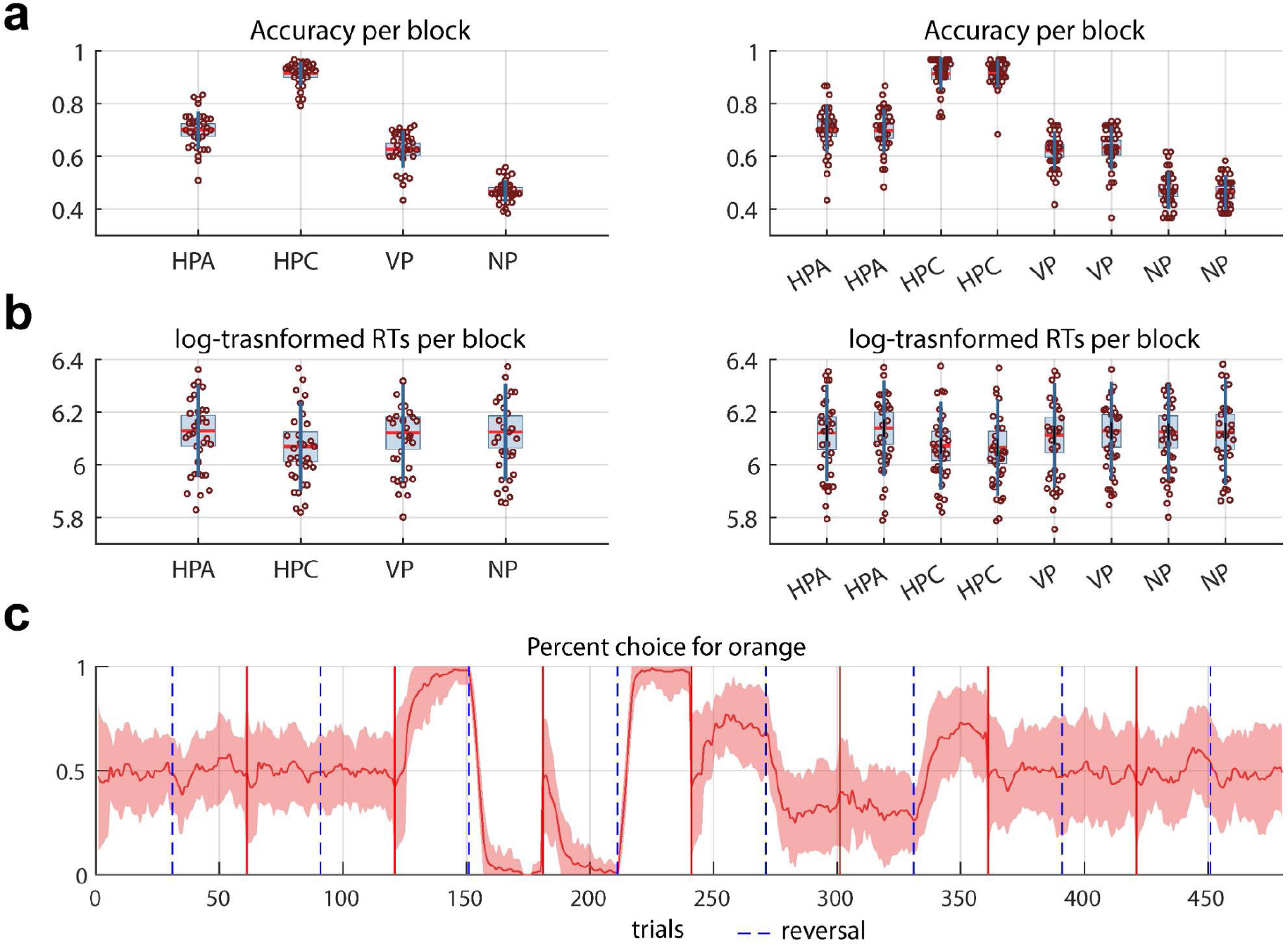
Main behavioural results. **(a)** Mean accuracy (number of time participants were rewarded) across the four main types of block (left panel) and across the four main types of blocks, played twice during one experimental session - HPA: highly-predictive anticorrelated block, HPC: highly-predictive correlated block, VP: variable predictive block, NO: non-predictive block. Left (**b)** Mean reaction times (log-transformed RTs) (number of time participants were rewarded) across the four main types of block (left panel) and across the four main types of blocks, played twice during one experimental session. **(c)** Percentage of choices for the orange colour across the eight blocks.

**Supplementary Figure 2.**
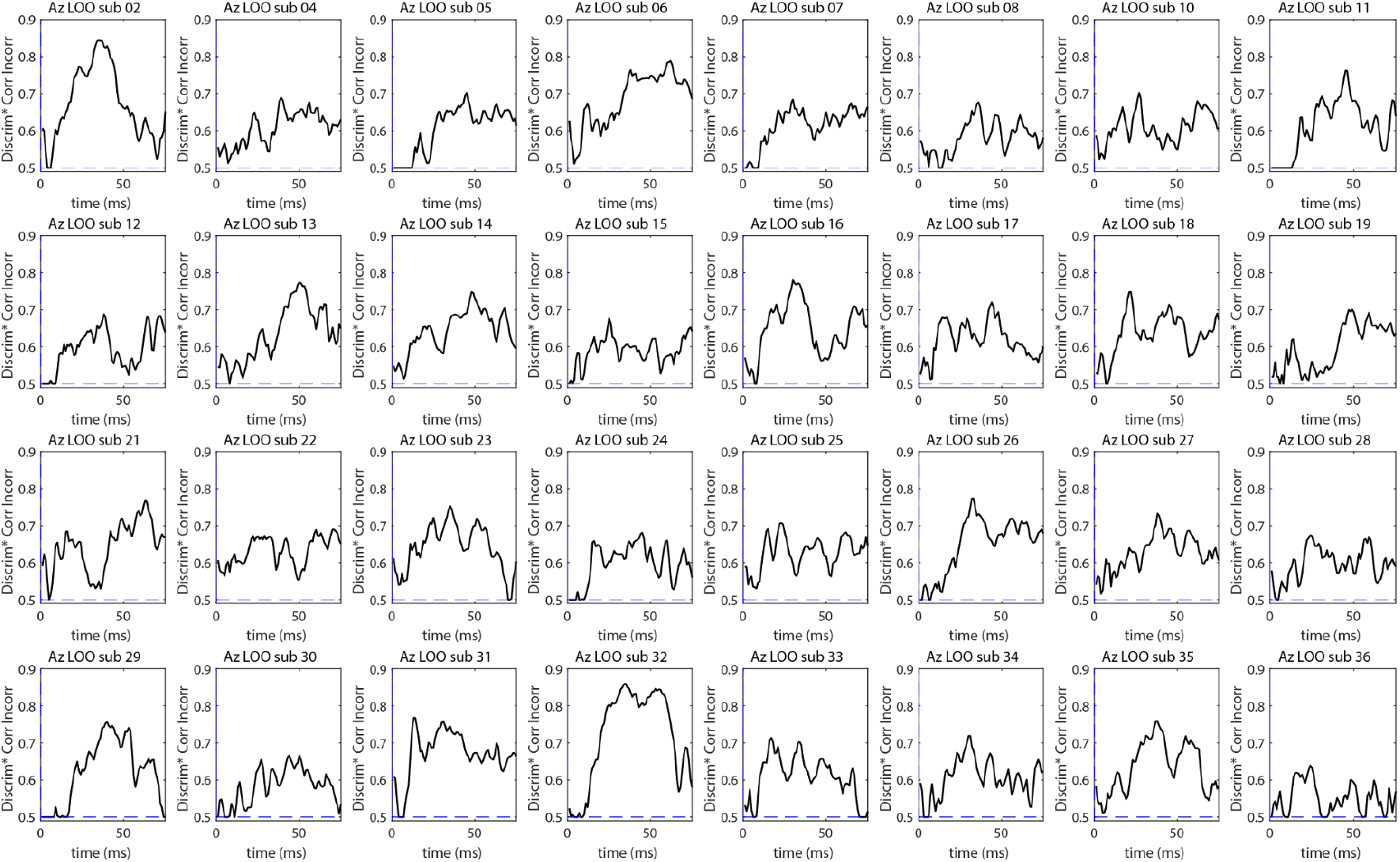
Discriminator performance (Az) during high-vs-low salient outcome discrimination of CRS-locked EEG data, for all subjects.

**Supplementary Figure 3.**
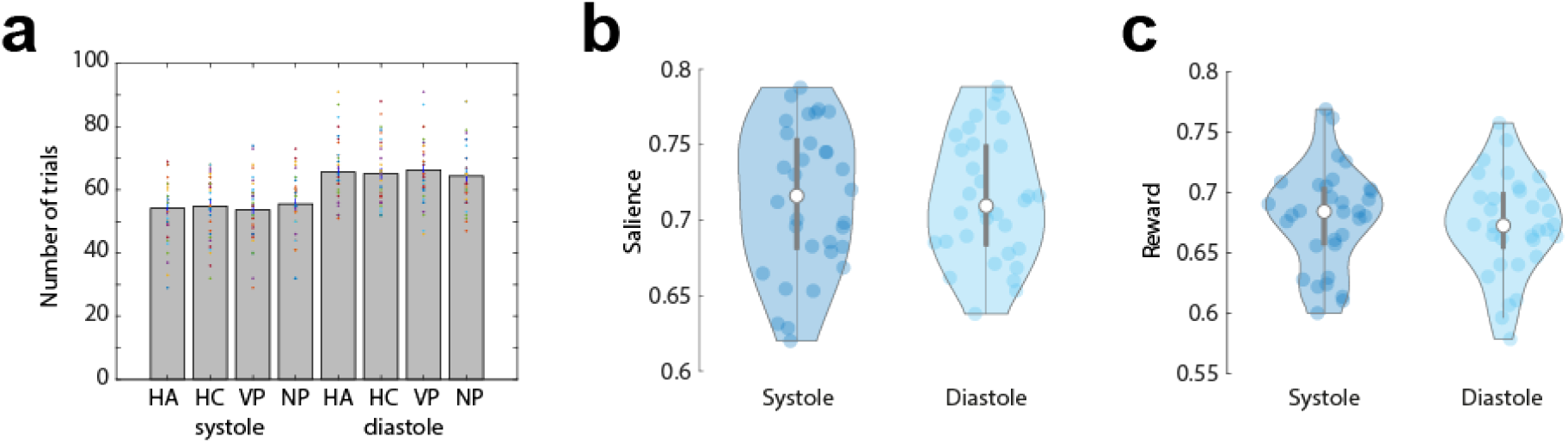
Control analyses for potential unaccounted effects. **(a)** The average number of trials for systole and diastole was the same across predictive and non-predictive blocks. **(b)** Mean absolute PE difference between the CRS for all outcomes presented at systole versus diastole (N=32). A violin plot is used to present all participants’ average. **(c)** Mean reward difference between the CRS for all outcomes presented at systole versus diastole (N=32). Neither the absolute PE (from the RL model) nor the reward was different between the systole and diastole outcomes.

**Supplementary Figure 4.**
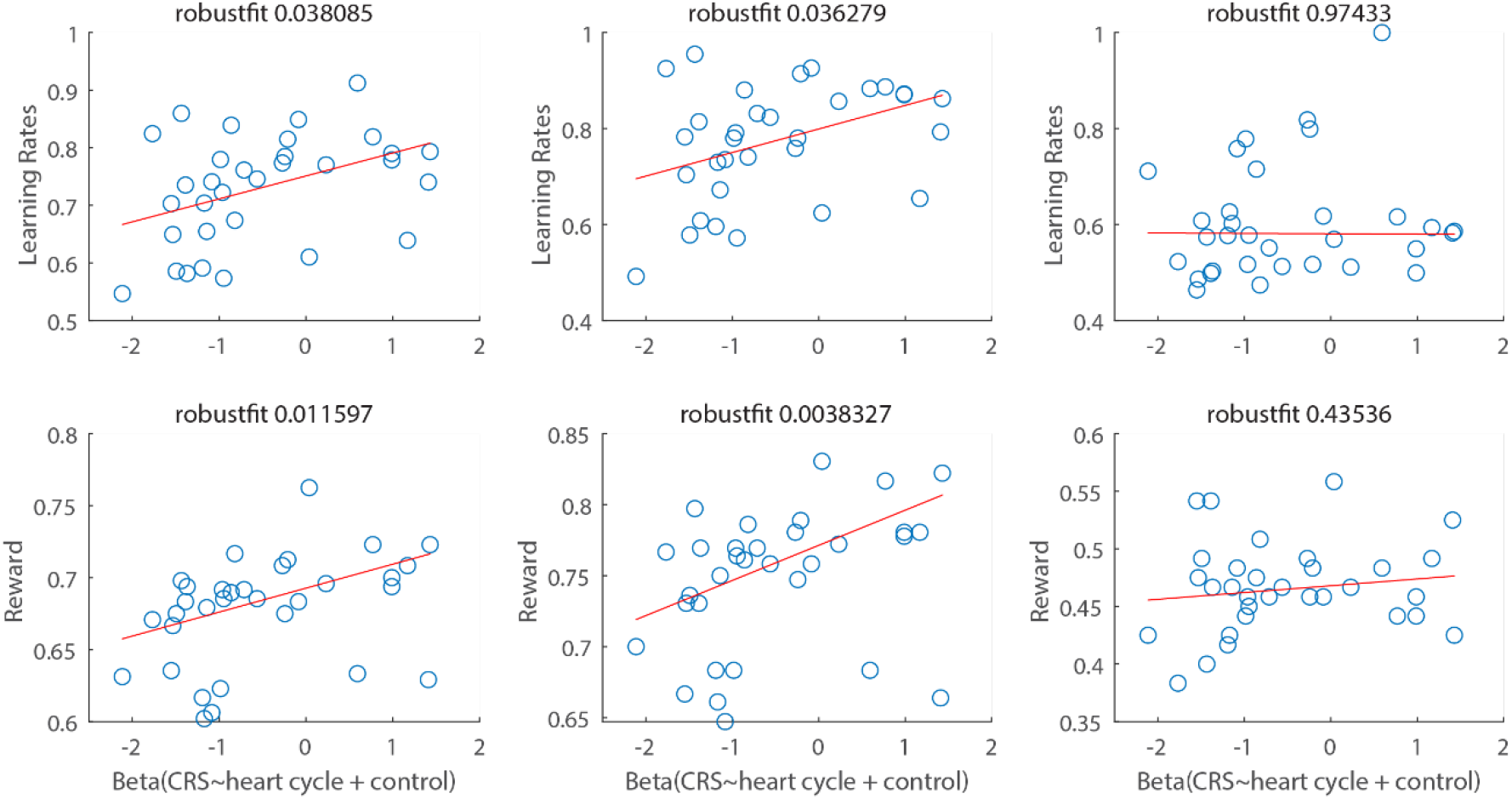
Control analyses for potential unaccounted effects of outcome (valence and absolute PE) in the relationship between CRS and heart cycle.

## Notes

### Competing Interest Statement

The authors have declared no competing interest.

